# From Sight to Insight: A Multi-task Approach with the Visual Language Decoding Model

**DOI:** 10.1101/2024.02.16.580578

**Authors:** Wei Huang, Pengfei Yang, Ying Tang, Fan Qin, Hengjiang Li, Diwei Wu, Wei Ren, Sizhuo Wang, Yuhao Zhao, Jing Wang, Haoxiang Liu, Jingpeng Li, Yucheng Zhu, Bo Zhou, Jingyuan Sun, Qiang Li, Kaiwen Cheng, Hongmei Yan, Huafu Chen

## Abstract

Visual neural decoding aims to unlock the mysteries of how the human brain interprets the visual world. While early studies made some progress in decoding visual activity for singular type of information, they failed to concurrently reveal the multi-level interweaving linguistic information in the brain. Here, we developed a novel Visual Language Decoding Model (VLDM) capable of decoding categories, semantic labels, and textual descriptions from visual perceptual activities simultaneously. We selected the large-scale NSD dataset to ensure the efficiency of the decoding model in joint training and evaluation across multiple tasks. For category decoding, we achieved the effective classification of 12 categories with an accuracy of nearly 70%, significantly surpassing the chance level. For label decoding, we attained the precise prediction of 80 specific semantic labels with a 16-fold improvement over the chance level. For text decoding, the scores of the decoded text surpassed the corresponding baseline levels by remarkable margins on six evaluation metrics. This study contributes significantly to extensive applications in multi-layered brain-computer interfaces, potentially leading to more natural and efficient human-computer interaction experiences.

## Introduction

The connection between vision and language has been a captivating area of research in cognitive neuroscience. Our brains exhibit intricate and close connections when processing visual and language information^1,2^. Studies indicate a close relationship between the processing of visual signals and the comprehension and production of language^3^. Visual signals are transmitted to the brain’s visual cortex^4,5^, while language processing primarily involves language-related brain areas like Broca’s and Wernicke’s areas^6,7^. However, there’s evidence of mutual interaction between visual and language brain areas^8^. When comprehending language, the brain activates regions associated with vision^9,10^, implying a potential interdependence between language comprehension and the processing of visual information. This collaboration might help explain why we use rich and vivid language expressions when faced with visual images or scenes.

The process of visual cognition, progressing from coarse to fine recognition^11,12^, is pivotal in perceiving and expressing what we see and feel about the world.. Taking the appreciation of an artwork as an example, depicted in Fig. 1, we express admiration for its beauty, but behind such verbal expression lies a gradual unfolding from the whole to the details. Initially, what we see is merely a categorization of the overall scene, such as determining whether it depicts a landscape, figures, or abstract art. As our appreciation proceeds, the visual system begins to process more specific semantic labels such as the sunset, lake, clouds, and birds, maybe coupled with some background details. Likewise, the description of the image may also undergo a hierarchical progression from the theme to the specifics. We might begin with category words like “scenery”, followed by foreground descriptive phrases such as “the lake illuminated by the glow of the setting sun”, “a bird freely soaring”, or “colorful clouds in the sky”. Our description might culminate in more vivid or poetic expressions such as “The setting sun and a solitary duck fly together, sharing the autumn water and the sky’s hue”. Therefore, the incremental process of language production mirrors the perception process in visual cognition^9,13^.

**Fig. 1.**
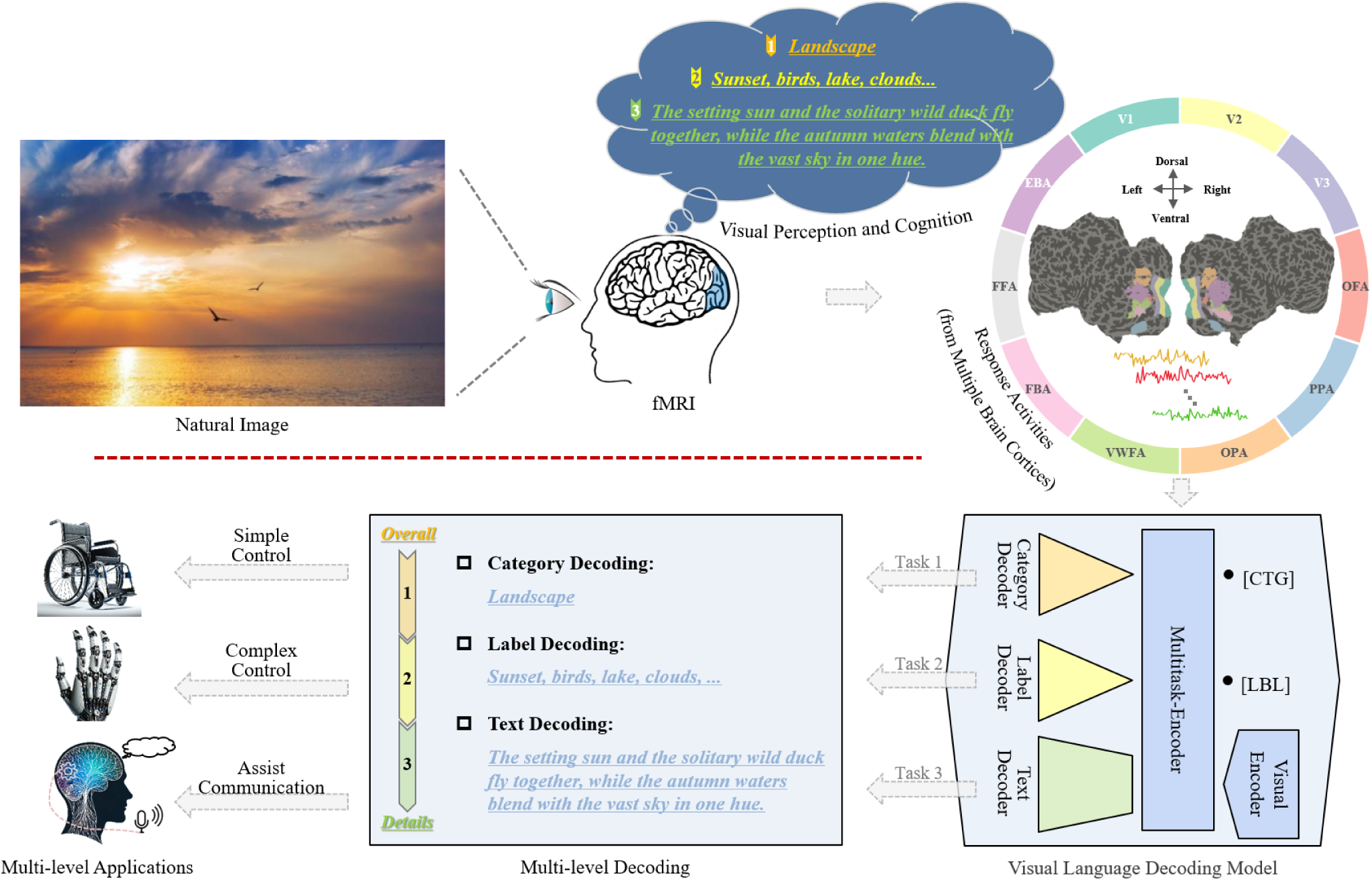
Overview of the Visual Language Decoding Process. Our study aims to decode simultaneously the primary categories, multiple labels, and textual descriptions from brain activity during the observation of natural images. Brain activity during the observation of natural images was captured using fMRI, covering multiple cortical regions. We selected 10 regions (V1, V2, V3, OFA, PPA, OPA, VWFA, FBA, FFA, and EBA) for conducting multitask decoding calculations. The Visual Language Decoding Model comprises the Visual-Encoder, Multitask-Encoder, Category-Decoder, Label-Decoder, and Text-Decoder. Initially, the Visual-Encoder and Multitask-Encoder encode the response activities from the chosen cortical regions to acquire multitask features of the stimulus images. Subsequently, the Category-Decoder, Label-Decoder, and Text-Decoder execute three individual decoding tasks simultaneously to predict the primary categories and multiple labels within the stimulus images, and to generate textual descriptions. The methods for these three decoding tasks correspond to different levels of brain-machine interface applications, namely, controlling wheelchairs, operating robotic arms, and restoring language functions.

How to decode linguistic information in visual cognition is an important research topic in brain science and artificial intelligence today. Since 2005, renowned labs worldwide have made significant progress in visual decoding using machine learning and deep learning techniques. These techniques are capable of extracting multi-level linguistic information including object categories^14,15^, semantic labels^16,17^ and descriptive text^18–20^ from visual activities triggered by natural scenes. However, these researches mainly decode information independently through a single task, which lacks knowledge sharing between different decoding tasks as well as limits the generalization capability of visual decoding. In the field of artificial intelligence, joint modeling of multiple tasks has been shown to help improve the efficiency of joint task processing. For example, in the field of computer vision, Wu et al. designed a multi-task network, YOLOP^21^, that is capable of handling three key driving perception tasks: object detection, drivable area segmentation, and lane detection, which greatly improves the performance of handling these three tasks simultaneously. Meanwhile, in natural language processing, cutting-edge technologies such as ChatGPT integrate GPT^22–24^, Reinforcement Learning from Human Feedback^25^, and Chain-of-Thought Prompt^26^ to accomplish advanced machine translation and Q&A systems. Inspired by technologies in computer vision and natural language processing, multi-task decoding methods will have the potential to improve the efficiency and generalization of visual information decoding.

Thus, we proposed a Visual Language Decoding Model (VLDM) which consisted of two encoders (Visual-Encoder and Multitask-Encoder) and three decoders (Category-Decoder, Label-Decoder, and Text-Decoder), capable of simultaneously performing three decoding tasks: primary categories, multiple labels, and textual descriptions. as illustrated in Fig. 1. Here our main objective is to achieve the synchronized decoding of visual perceptual activities into multi-level information. The responses to visual activities are acquired through functional magnetic resonance imaging (fMRI), measuring blood-oxygen-level dependent (BOLD) signals. fMRI offers high spatial resolution and is a commonly used non-invasive measurement tool^27^. We selected the data of four subjects (Sub 1, Sub 2, Sub 5, and Sub 7) from the large publicly available NSD dataset^28^ and collected responses from 10 visual cortex regions while they viewed 30,000 natural images from the COCO dataset^29^ which provided 12 primary categories, 80 multiple labels, and5 textual descriptions for each natural image. Hopefully, our proposed Visual Language Decoding Model (VLDM) will take advantage of the well-suited dataset to predict multiple levels of language information by integrating three decoding task which correspond to at least three levels of application value for brain-machine interfaces: 1) Category decoding aids in controlling assistive devices like wheelchairs via brain activity, enhancing daily life convenience for disabled individuals; 2) Label decoding provides more complex and advanced control capabilities for disabled individuals, such as operating robotic arms, promoting more flexible lifestyles; 3) Text decoding supports aphasic patients in restoring language function, enabling natural communication and instruction transmission.

## Results

In this section, we will sequentially present the relevant research findings concerning category decoding, label decoding, and text decoding tasks.

### Category decoding results

We employed classification accuracy as an evaluation metric, quantitatively analyzing the accuracy of correctly categorizing viewed natural images into the 12 distinct categories (person, vehicle, outdoor, animal, accessory, sports, kitchen, food, furniture, electronic, appliance, and indoor) (depicted in Fig. 2a). During the training process of the multitask decoding model, we meticulously recorded the classification accuracy of four subjects on the test set after each iteration, as depicted in Fig. 2b. The results indicate a noticeable consistency in the accuracy variation curves among the four subjects. Upon reaching a stable state after a gradual increase in the early iterations, the classification accuracy for the four subjects respectively reached 0.6548, 0.6413, 0.6761, and 0.6099, significantly surpassing the chance level (1/12, 0.083).

**Fig. 2.**
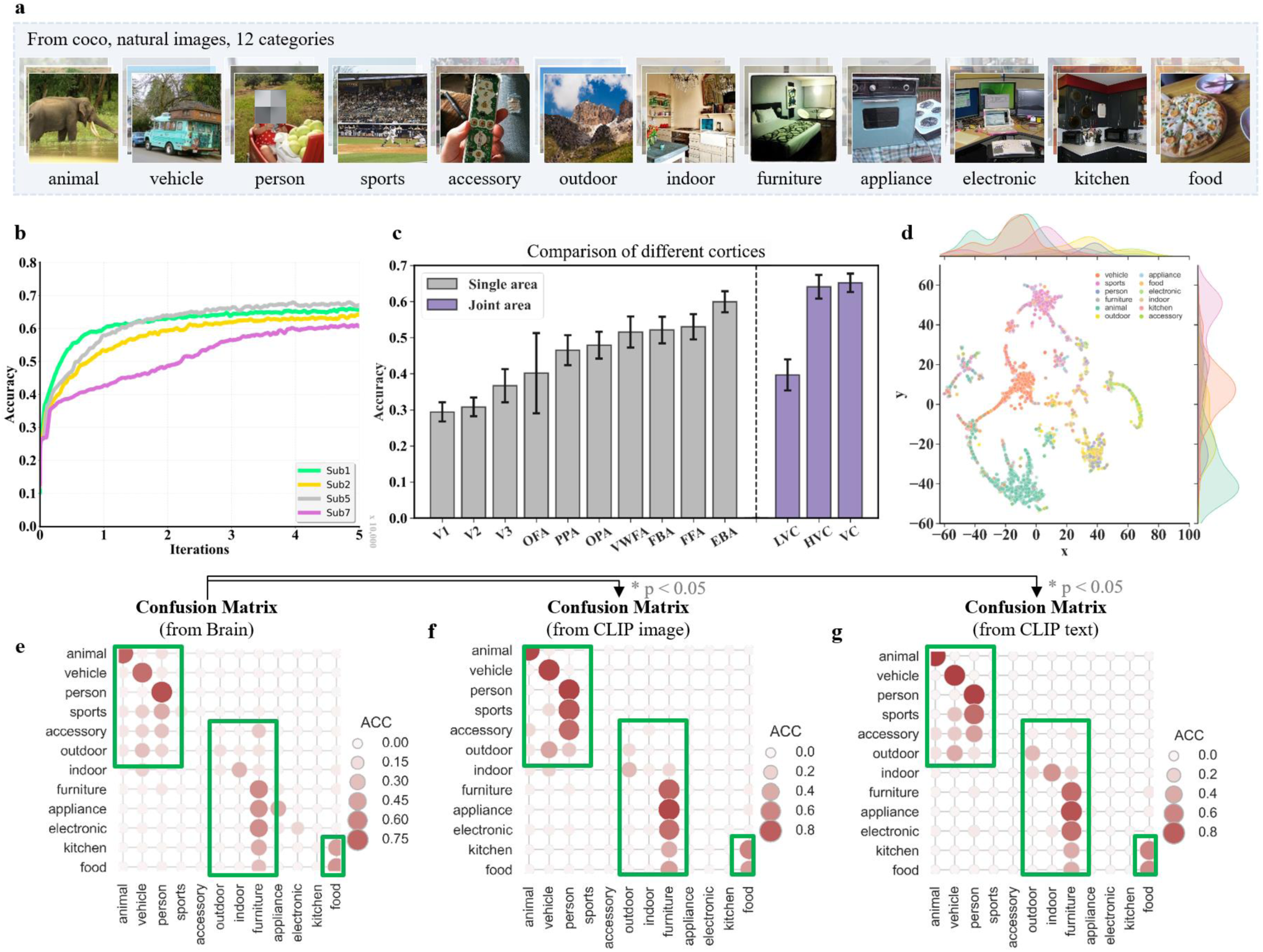
Illustrates the results of category decoding. **a**, displays sample images from twelve principal classes sourced from the COCO dataset. **b**, presents the accuracy trendlines from the testing set of four subjects throughout the training phase. **c**, compares the classification accuracies of different cortical regions for the 12 natural image categories. The error line indicates the standard deviation of accuracy for the four subjects. Gray and purple bars indicate the average accuracy rates produced by the Single and Joint visual areas. **d**, shows the final layer output features of the category decoder, which are downscaled to 2D space by T-SNE to obtain visualization results. Each point represents each sample of the test set, and different colors indicate different categories. **e**, confusion matrix for 12 categories obtained from human brain activity. **f**, confusion matrix for 12 categories obtained from CLIP image features. **g**, confusion matrix for 12 categories obtained from CLIP text features.

### (1) Comparison of classification accuracy across different visual cortical areas

In Fig. 2c, the results reveal noticeable differences in the classification accuracy among various visual cortical regions. Among the independent visual areas, the EBA region exhibited the highest classification accuracy in visual decoding tasks, reaching 0.5994. Following EBA, FFA, FBA, and VWFA regions achieved accuracies of 0.5302, 0.5210, and 0.5153, respectively. The lower-level visual cortex areas, V1, V2, and V3, showed relatively lower classification accuracies at 0.2940, 0.3079, and 0.3668, respectively. OPA and PPA regions had classification accuracies of 0.4792 and 0.4653, respectively, while OFA demonstrated a slightly lower accuracy at 0.4014. VC, the combined visual areas, demonstrated the highest classification accuracy at 0.6520, followed closely by HVC at 0.6409, while LVC exhibited an accuracy of 0.3968. Taken together, it’s evident that higher-level visual areas (such as EBA, FBA, FFA, OFA, OPA, PPA, and VWFA) typically exhibit stronger classification decoding abilities than the lower-level visual areas (e.g., V1, V2, and V3) and the combined visual areas (LVC, HVC and VC) had higher classification accuracies than any of their constituent visual areas.

### (2) 2D Visualization of output features from the classification decoder

Using the t-distributed Stochastic Neighbor Embedding (t-SNE) method, we reduced the category distribution features to two-dimensional(2D) space. In this 2D space, closer distances indicate higher correlation between category distribution features, while farther distances signify lower correlation. Fig. 2d showcases the t-SNE visualization results for Subject 1. The results illustrate that the sports category and animal category are relatively positioned on opposite sides, reflecting their distinctiveness in the feature space and effectively demonstrating the efficacy of the multitask decoding model in recognizing dissimilar categories. Similarly, close proximity between the kitchen and food categories is observed, representing objects associated with similar real-world environments, such as bread commonly being found in kitchens. Moreover, the vehicle category is positioned between the sports and outdoor categories, mirroring the fact that vehicles like motorcycles or bicycles are often associated with human movement and outdoor settings in real-world scenarios. These visualizations “coincidentally” reproduce the real-world relationships among these categories, thereby validating the effectiveness and accuracy of our classification decoder. Additionally, Supplementary Fig. 1 displays the t-SNE visualizations for Subject 2, Subject 5, and Subject 7.

### (3) Confusion matrix for classification

Fig. 2e displays the average confusion matrix across the four subjects. Each cell in the confusion matrix shows the frequency of a specific category being misidentified by the decoder as another category. Specifically, the rows of the matrix represent the actual categories, while the columns represent the predicted categories. Therefore, the “size of the circles” in each cell indicates the frequency of instances where the actual category was predicted as a particular category. Ideally, most instances should be concentrated along the diagonal of the confusion matrix, signifying agreement between predicted and actual categories. In Fig. 2e, it’s noticeable that there are larger “red circles” along the diagonal, indicating these categories are correctly classified, reflecting the overall accuracy of our model. For instance, categories like animal, vehicle, person, furniture, and food are easily correctly classified. However, there are also instances of misclassifications. For example, the sports category is most frequently misclassified as the person category; accessory, outdoor, and indoor categories are commonly misclassified as the vehicle category; appliance, electronic, and kitchen categories are frequently misclassified as the furniture category; and the kitchen category is most prone to being misclassified as the food category. We conducted two additional experiments: 1) classifying CLIP features of natural images to obtain a confusion matrix for the 12 categories (as shown in Fig. 2f); 2) classifying CLIP features of textual descriptions of natural images to obtain a confusion matrix for the 12 categories (as shown in Fig. 2g). It was discovered that both the confusion matrices derived from natural images and textual descriptions exhibited very high similarity to the confusion matrix derived from brain data. Additionally, Supplementary Fig. 2 demonstrates the confusion matrices for subjects 1, 2, 5, and 7.

### Label decoding results

The natural images involved in this study encompass a total of 80 different labels (refer to Supplementary Table 1 for details). It’s noteworthy that each natural image contains only a few labels, as depicted in Fig. 3a. In Fig. 3b, it is observed that the accuracy curve for label decoding exhibits an upward trend as the training progresses. When the accuracy curve reaches a stable state after a certain number of iterations, the accuracy rates for the four subjects are 0.1940, 0.1811, 0.2123, and 0.1529, respectively, with the average accuracy (0.1851) being approximately 15 times higher than the chance level (0.0125). It indicates that the performance of the multitask decoding model in label decoding significantly surpasses random guessing. In Fig. 3c, the results indicate varied performance in decoding different labels. Some labels such as “person”, “clock”, and “sink”, occupying relatively higher proportions in visual responses at 18.12%, 1.57%, and 1.56% respectively, exhibited generally higher average decoding accuracies of 0.9192, 0.4171, and 0.4140. However, for less frequently occurring labels like “toaster” (0.07%), “hair drier” (0.06%), and “parking meter” (0.21%), the model displayed relatively lower decoding accuracies at 0.0032, 0.0045, and 0.0047 respectively. Nonetheless, except for these three less frequent labels, most other labels displayed accuracies significantly surpassing the chance level.

**Fig. 3.**
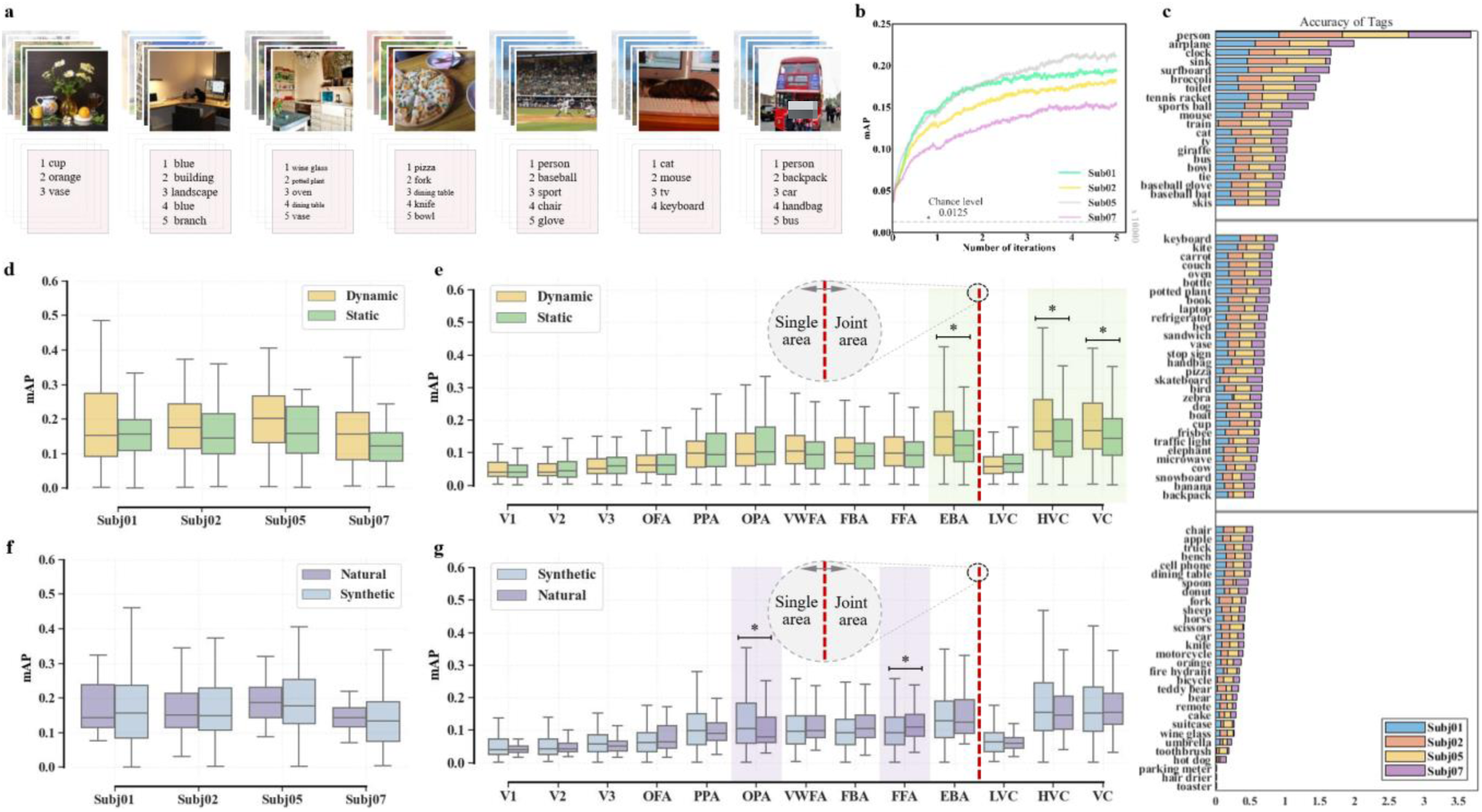
depicts the research findings on label decoding. **a**, shows a visualization of the label decoding results, where the upper section displays images from the test set stimuli, while the lower section represents various labels decoded from brain visual activity. **b**, describes the variation curve of label accuracy obtained from brain visual activity based on four test subjects, with the dashed line indicating the level of random prediction. **c**, presents the accuracy rates of the four test subjects across 80 labels. **d**, displays the comparison of accuracy between “Moving attributes” and “Static attributes” labels. **e**, provides a comparative analysis of the accuracy of “Dynamical attributes” and “Static attributes” labels across 10 independent visual cortices and 3 combined visual cortices. **f**, illustrates the comparison of accuracy between “Natural attributes” and “Synthetic attributes” labels. **g**, conducts a comparative analysis of the accuracy of “Natural attributes” and “Synthetic attributes” labels across 10 independent visual cortices and 3 combined visual cortices.

### (1) Comparison of accuracies between moving attribute and static attribute labels

In Fig. 3d, we compared the decoding accuracy of these two types of attribute labels. The results indicated that, decoding accuracy was higher on average for moving labels (0.1923) compared with static attribute labels (0.1594), but both surpassed the random baseline of 0.0125. Next, we further explored the distribution of “moving attributes” and “static attributes” in the visual cortex. In Fig. 3e, we compared the decoding accuracy of these two types of attribute labels across 10 distinct visual areas (V1, V2, V3, EBA, FBA, FFA, OFA, OPA, PPA, and VWFA) and 3 combined visual areas (LVC, HVC, and VC). When observing the independent visual areas, we found a notably higher decoding accuracy for moving attribute labels than for static attributes in the EBA region. This primarily indicates the specific sensitivity of the EBA region towards moving attributes when processing visual information.

### (2) Comparison of accuracy between natural and artificial attribute labels

Meanwhile, to explore the decoding performance differences between “natural attributes” and “artificial attributes”, we categorized 80 labels into these two groups. As depicted in Fig. 3f and Fig. 3g, this is an intriguing finding that the OPA and FFA regions might exhibit slight preferences or advantages when processing “natural attributes” or “artificial attributes”. Specifically, the OPA region shows slightly higher decoding accuracy when processing “natural attributes” and relatively lower accuracy when dealing with “artificial attributes”. Conversely, the FFA region demonstrates the opposite pattern.

### Text decoding results

Fig. 4a shows the results of text decoding, where each block’s image represents a natural image viewed by the subject, and the text represents the sentence generated by our proposed decoding model. For example, in Fig. 4a, the decoded sentence for the second image in the first row is “a cat is sitting on a wooden chair”, where “cat” and “wooden chair” accurately describe the objects in the image, while “sitting” accurately reflects the action of the objects in the image, and prepositions and articles string these objects and actions into a descriptive sentence. These results indicate that our model can accurately capture not only the main objects and actions in the image but also effectively organize these semantic labels into a complete and coherent descriptive text. For further insights into the results of text decoding, please refer to Supplementary Fig. 3 and Supplementary Fig. 4.

**Fig. 4.**
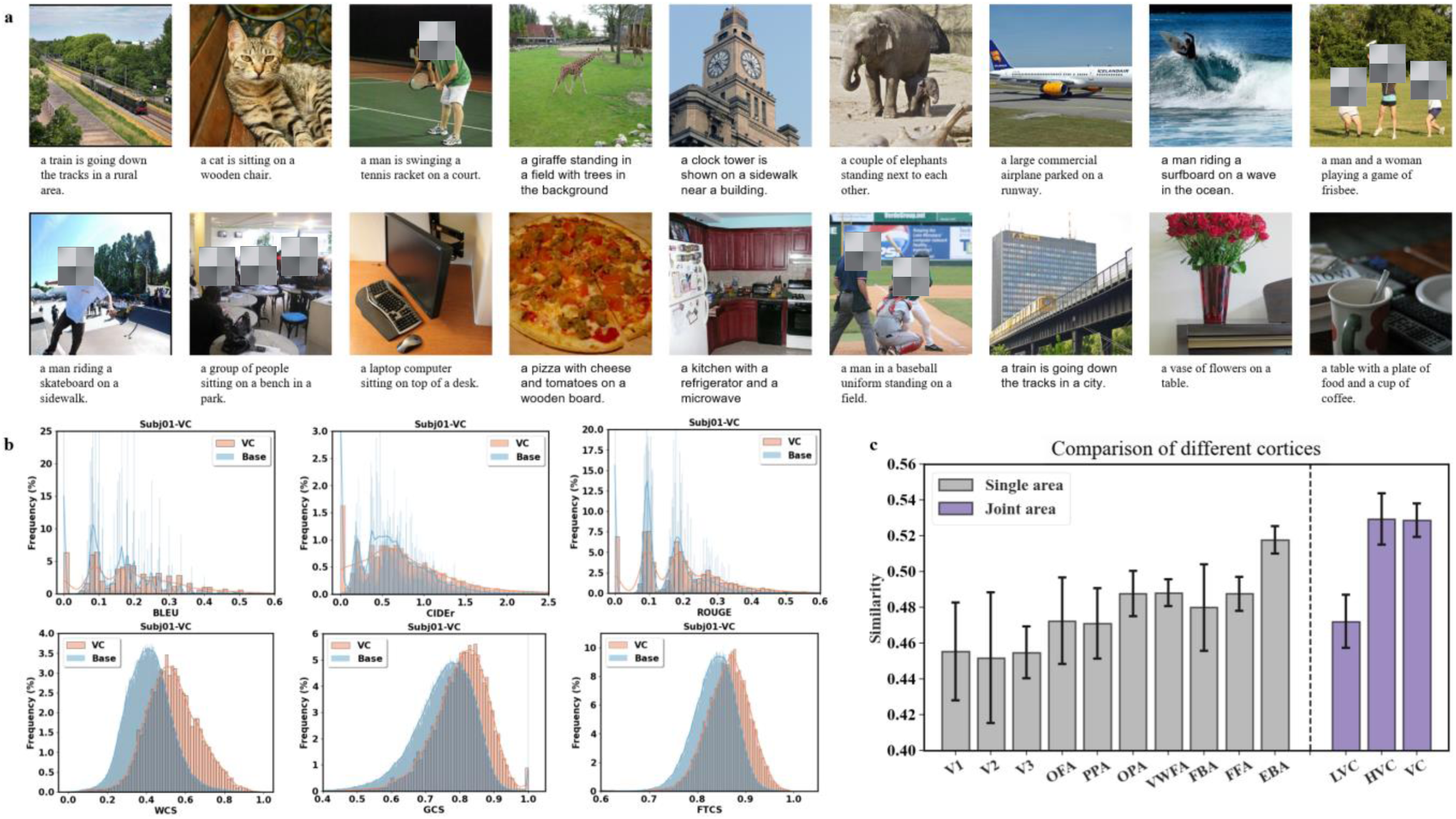
illustrates the results of text decoding. **a**, displays a visual representation of the language decoding results: the upper section exhibits stimulating images from the test set, while the lower section presents sentences derived from brain visual activity describing the stimulating images. **b**, reveals histograms depicting the frequency distribution of six evaluation metrics. In each subplot, the blue distribution represents evaluation scores between decoded text and standard text from the test set, while the orange distribution serves as a baseline distribution, showcasing evaluation scores between decoded text and all text from the test set. **c**, showcases a comparison of language decoding accuracy across ten independent visual cortices and three combined visual cortices, where the dashed line represents the level of random prediction.

### (1) Accuracy assessment of text decoding

On the test set, the average BLEU, CIDEr, and ROUGE scores across four subjects were 0.1922, 0.7986, and 0.1909, respectively. Their corresponding baseline levels were 0.1421, 0.5843, and 0.1439, respectively. These baseline levels referred to the average scores of the decoded sentences compared to all sentences in the test set across the respective evaluation metrics. Relative to the baseline levels, the BLEU, CIDEr, and ROUGE scores increased by over 35%, 36%, and 32%, respectively. Furthermore, compared to the baseline levels, the WCS, GCS, and FTCS scores (0.5287, 0.8020, and 0.8612) increased by over 28%, 6%, and 3%, respectively. Fig. 4b displays histograms of the six evaluation metrics obtained in the VC region for subject 1. In each subplot, the yellow distribution represents the scores of the respective evaluation metrics between the decoded sentences and the target sentences in the test set, while the blue distribution represents the scores between the decoded sentences and all sentences. Additionally, Supplementary Fig. 5 illustrates histograms of six evaluation metrics obtained in the VC region for subjects 2, 5, and 7. The yellow distribution generally leans towards the right side, indicating that the sentences generated by our model are semantically closer to the target sentences.

### (2) Comparison of text decoding accuracy in different visual cortices

Fig. 4c vividly illustrates the significant differences in decoding accuracy across various visual regions. Among them, EBA exhibited the highest average score at 0.5178, demonstrating the most optimal decoding accuracy. Following this, OPA, VWFA, and FFA had decoding accuracies of 0.4876, 0.4881, and 0.4876; FBA, OFA, and PPA scored 0.4799, 0.4725, and 0.4711; V1, V2, and V3 displayed the lowest decoding accuracies at 0.4552, 0.4518, and 0.4547, respectively. Meanwhile, the combined regions, HVC and VC, showed very similar and notably higher decoding accuracies (0.5293 and 0.5287) while LVC exhibited a slightly lower accuracy of 0.4722. In summary, we found that the decoding accuracies in combined regions were slightly higher than those in individual areas: the accuracy of LVC surpassed that of V1, V2, or V3; the accuracy of HVC also exceeded any of the ten independent regions.

## Discussion

This study established a multi-task joint decoding model called VLDM to extract multi-level language information from the brain’s visual cortex. By utilizing NSD and VLDM, we have explored how the brain presents different levels of language symbols when processing visual information, including category, semantic labels, and text descriptions.

First of all, the results of the category decoding revealed a gradual improvement up to a stabilization in classification accuracy across four subjects as the number of iterations increased. This trend potentially supports the robustness of the model, indicating a relative consistency in its performance across different individuals. Similar tendencies were observed in our previous studies although specific accuracy values varied^15^. These differences might stem from various factors such as the datasets used, the selection of visual cortical regions, and the design of decoding models. Furthermore, a comparative analysis of classification accuracy demonstrated significant differences among distinct visual cortical regions. Individual higher-order visual areas exhibited higher accuracy while lower-level visual cortical areas displayed relatively lower accuracy. These outcomes align with previous research supporting the viewpoint that higher-level visual areas often demonstrate stronger classification decoding capabilities^15,30^. Concerning combined visual regions, the LVC region demonstrated higher classification accuracy than any of V1, V2, and V3, while HVC displayed higher accuracy than any of its constituent regions. These findings support the idea that integrating multiple visual regions can enhance decoding performance^31^. Remarkably, this study employed t-SNE analysis to visualize the category distribution features outputted by the category-decoder in a 2D format, revealing the interrelations among different categories in real world. For instance, the far distance between ‘sports’ and ‘animals’ reflected their distinct feature spaces, while ‘kitchen’ and ‘food’ categories were closely clustered, consistent with the real-world scenario where objects representing these concepts often coexist in similar environments. Similar observations have been noted in previous research^32,33^. Finally, the confusion matrices of classification showed a high similarity between image or text and brain activity. This extends previous research, suggesting the patterns of confusion between categories, whether in natural images or language descriptions, can be reflected in the neural visual activity of the brain. It provides detailed information for specific categories and enables a deeper investigation of misjudged categories, allowing for fine-tuning of the model to improve decoding performance.

Second, we found variations in the decoding accuracies among different labels. Higher-frequency labels, such as “person”, “clock”, and “sink”, exhibited relatively higher average decoding accuracy, whereas lower-frequency labels like “toaster”, “hair drier”, and “parking meter” showed relatively lower decoding accuracy. This contrast might stem from low frequencies of these labels in the training samples, thereby limiting the model’s learning and adjusting performance. This divergence aligns with similar phenomena observed in prior research that labels appearing less frequently in training data may pose greater challenges to models due to their limited sample count, making it harder to adequately learn their features and patterns^34,35^. This underscores the model’s high accuracies across the majority of labels, demonstrating its adaptability and robustness for multi-label decoding tasks.

Moreover, the comparison between moving and static attributes deepened our understanding of semantic attribute decoding in the brain. Earlier studies have highlighted differences in how distinct brain regions process static and dynamic information^36^. This outcome reveals the specificity of EBA in processing dynamic information, consistent with previous evidence that the EBA region excels in handling information related to body movements or object motion^37,38^. One possible explanation is that moving attribute labels involve richer and more complex information, such as motion direction, speed, object changes, etc. Meanwhile, the study delved into the preferences for natural and man-made attribute labels within specific brain regions. The OPA was found to have a certain preference for man-made attribute labels, whereas FFA exhibited a preference for natural attribute labels. This disparity might arise from the specialized visual cognitive functions of these two regions. The OPA region is believed to primarily involve object recognition and localization in complex scenes^39^, and in modern human life, most encountered objects are human-made, such as phones, computers, cars, etc. In contrast, the FFA region is a crucial area for facial recognition^40^, showing higher decoding accuracy when processing “natural attributes” like faces. These results underscore the functional specificity of different brain regions in processing visual information, providing clues for understanding how the brain concurrently processes and integrates various kinds of visual information. However, further research is needed to validate the repeatability of these observations regarding decoding these pairs of attributes and to delve deeper into the neural mechanisms underlying these differences.

Third, our multitask visual-language decoding model succeeded in accurately capturing key objects and actions within images, organizing them into fluent and comprehensible textual descriptions. This echoes well with our prior work which directly utilized a dual-channel language model to generate precise language information from fMRI neural activity^20^. Furthermore, our findings indirectly corroborated that higher brain areas, such as FFA and VWFA, are primarily responsible for handling high-level semantic information, while lower-level brain regions like V1 and V2 handle fundamental visual information^41–43^. Previous research has indicated the pronounced advantages of FFA and VWFA areas in processing information related to faces and bodies^43^. Meanwhile, the involvement of HVC in recognizing semantic objects and scenes further supports the greater contribution of these regions to language decoding than those lower areas focusing on basic visual features^14^. This intrinsic mechanism of complementary collaboration between vision and language in the brain warrants further exploration in subsequent studies.

Taken together, this study innovatively decodes categories, labels, and textual descriptions simultaneously, addressing previous gaps in exploring how the brain intertwines and interacts with various visual information. This research emphasizes the complexity of brain processing of visual information, contributing to a more comprehensive understanding of the interplay of visual cognition and language processing within the brain. Exploring visual information processing mechanisms from the perspective of brain decoding may offer new insights for the development of brain-computer interface technology and future human-computer interactions.

However, there are some limitations. The NSD dataset might limit the model’s generalizability in terms of scale and diversity. Next, the scale of the decoding model might influence its performance. Future research could consider utilizing larger and more diverse datasets to enhance the model’s generalization capabilities across various scenes, objects, and contexts. Additionally, there is a need to work on the interpretability of the model to comprehensively understand the decision-making mechanisms during both decoding and predicting processes. Future research should also explore universal decoding techniques akin to ChatGPT, paving the way for new frontiers in brain decoding research.

## Methods

### The NSD and the COCO Dataset

The natural scene dataset NSD^28^ is a large-scale fMRI dataset comprising data from 8 subjects. Each subject viewed nearly 10,000 distinct colored natural scene images, each seen thrice, totaling 30-40 scanning sessions. We selected data from 4 subjects (Sub 1, Sub 2, Sub 5, and Sub 7). This data underwent two preprocessing steps involving temporal and spatial interpolation to correct slice timing differences and head motion. For decoding tasks, we utilized Z-score single-trial beta values generated by the GLMSingle method. The NSD dataset delineates multiple regions of interest (ROIs). We chose 10 distinct visual areas: V1, V2, V3, OFA, PPA, OPA, VWFA, FBA, FFA, and EBA. Additionally, we defined three combined regions: LVC, HVC, and VC. Specifically, LVC represented the combined region of V1, V2, and V3; HVC represented the combined region of OFA, PPA, OPA, VWFA, FBA, FFA, and EBA; whereas VC denoted the overall region encompassing these 10 distinct visual areas. The dimensions of these regions’ response activities were standardized (up-sampled or down-sampled) to the same length (1024 dimensions), with their original dimensions (also referred to as voxel count) detailed in Table 1.

**Table 1.** The original dimensions (Number of voxels) of visual activities from different ROIs in NSD.

|  | V1 | V2 | V3 | EBA | FBA | FFA | OFA | OPA | PPA | VWFA |
| --- | --- | --- | --- | --- | --- | --- | --- | --- | --- | --- |
| Sub 1 | 1350 | 1433 | 1187 | 2971 | 826 | 794 | 355 | 1611 | 1033 | 1278 |
| Sub 2 | 1102 | 1075 | 1097 | 3439 | 1217 | 869 | 441 | 1381 | 994 | 552 |
| Sub 5 | 1113 | 1081 | 925 | 4587 | 968 | 907 | 782 | 1332 | 1221 | 697 |
| Sub 7 | 1142 | 986 | 726 | 3062 | 552 | 484 | 316 | 1083 | 912 | 1068 |

The stimulus images in the NSD were derived from the COCO dataset^29^. Each subject completed 40 scanning sessions, with each session comprising 750 stimulus images. In this paper, we selected the first 37 sessions, totaling 27,750 stimulus images. Among these, 2,770 stimulus images were used as a test set, while the remaining 24,980 were used as a training set. Each stimulus image included a primary category, multiple labels, and five text descriptions. The primary categories, termed as “super-category” in the COCO dataset, encompassed a total of 12 categories: person, vehicle, outdoor, animal, accessory, sports, kitchen, food, furniture, electronic, appliance, and indoor. The multiple labels, termed as “name” in the COCO dataset, amount to 80 labels. The distribution of samples for these primary categories and labels is detailed in Supplementary Table 1. The five text descriptions originate from natural image descriptions in the COCO dataset, and during subsequent model training, we randomly selected one sentence for participation in the training. In summary, for each subject, we obtained datasets consisting of 24,980 training samples and 2,770 test samples. Each sample comprised a natural image, a primary category, multiple labels, five text descriptions, as well as visual activities from ten distinct regions and three combined regions.

### The Encoder-Decoder-Based VLDM

To investigate the multi-layered decoding of visual perception, we developed a VLDM which consists of two encoders (Visual-Encoder and Multitask-Encoder) and three decoders (Category-Decoder, Label-Decoder, and Text-Decoder). These encoders and decoders play distinct roles in the process of decoding visual information. Fig. 5 illustrates the overall structure of VLDM.

**Fig. 5.**
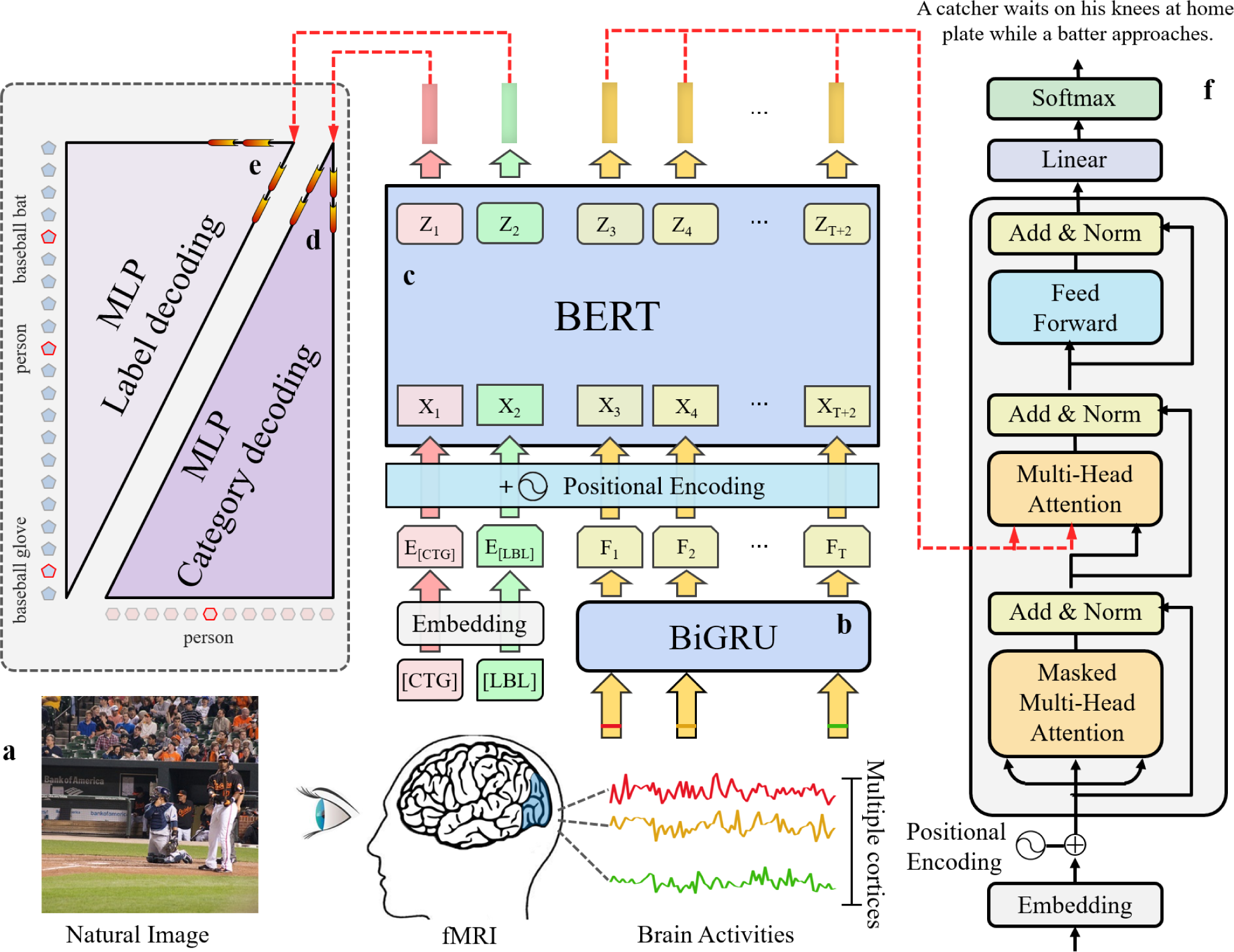
illustrates the encoding-decoding framework of the visual language decoding model. **a**, overview of the natural image perception experiment: Subjects view natural images while their brain activity is captured via fMRI. Subsequently, visual response activities are extracted from multiple brain cortices for subsequent information decoding tasks. **b**, Visual-Encoder: Constructed using BiGRU, it aims to receive visual response activities from multiple brain cortices, process them, and output latent features of visual activity. **c**, Multitask-Encoder: Built with BERT, its objective is to receive latent features of visual activity, integrate two tokens for category and label, and further process to output multitask features. **d**, Category-Decoder: Constructed with MLP, its function is to predict the primary category of stimulus images based on the [Z_1_] feature within multitask features. **e**, Label-Decoder: Also constructed with MLP, its function is to predict multiple labels of stimulus images based on the multitask feature [Z_2_]. **f**, Text-Decoder: Constructed using Transformer decoder, its function is to generate textual descriptions of stimulus images based on multitask features [Z_3_, Z_4_, . . ., Z_T+2_]. **Note: BiGRU stands for Bidirectional Gated Recurrent Unit; BERT stands for Bidirectional Encoder Representation from Transformers; MLP stands for Multi-layer Perceptron.

### (1) Visual-Encoder

While viewing natural images (visual stimuli), subject’s BOLD responses (brain activities) were measured using fMRI, as depicted in Fig. 5a. The design of the Visual-Encoder is to capture semantic information about the stimulus image from perceptual activities. We use a Bidirectional Gated Recurrent Unit (BiGRU) as the architecture for the Visual-Encoder. Illustrated in Fig. 5b, the Visual-Encoder initially receives visual activities (V_1_, V_2_, . . ., V_T_) from multiple brain cortices (ROIs), subsequently processing and obtaining visual features (F_1_, F_2_, . . ., F_T_), where T represents the selected number of brain cortices. Here, we selected 10 brain cortices, comprising: V1, V2, V3, OFA, PPA, OPA, VWFA, FBA, FFA, and EBA. These visual activities are fed into the BiGRU^44^, Bidirectional Gated Recurrent Unit, to acquire visual features (F_1_, F_2_, . . ., F_T_).

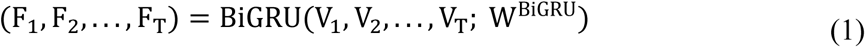

where W^BiGRU^ denotes the set of parameters included in BiGRU.

Subsequently, these visual features (F_1_, F_2_, . . ., F_T_) carrying semantic information are used as the main body of the input for the Multitask-Encoder.

### (2) Multitask-Encoder

The goal of the Multitask-Encoder is to integrate [CTG] and [LBL] tokens upon the foundation of visual features (F_1_, F_2_, . . ., F_T_) to obtain the multitask features (Z_1_, Z_2_, . . ., Z_T+2_), as depicted in Fig. 5c. [CTG] and [LBL] correspond respectively to category and label decoding tasks. To facilitate diverse task execution, we concatenate the two embedding vectors associated with [CTG] and [LBL], along with the visual features (F_1_, F_2_, . . ., F_T_) obtained from the Visual-Encoder. Moreover, to enable the model to learn positional dependencies within the sequence, we introduce positional embeddings^45^, creating visual features (X_1_, X_2_, . . ., X_T+2_) enriched with positional encoding. Subsequently, to capture advanced visual information, we utilize BERT^46^ as the architecture for the multi-task encoder. BERT receives visual features (X_1_, X_2_, . . ., X_T+2_) with positional encoding for computing and outputting multitask features (Z_1_, Z_2_, . . ., Z_T+2_). The specific formula is articulated as follows:

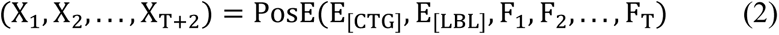

where E_[CTG]_ and E_[LBL]_ denote the embedding vectors corresponding to [CTG] and [LBL], respectively; and PosE(⋅) denotes the position processor.

After the collaborative work of the Visual-Encoder and Multitask-Encoder, the visual activities (V_1_, V_2_, . . ., V_T_) are encoded into multitask features (Z_1_, Z_2_, . . ., Z_T+2_). The specific formula is as follows:

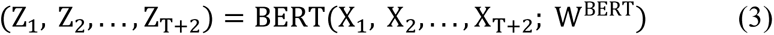

where BERT(⋅) and W^BERT^ denote BERT and its parameter set, respectively.

Afterwards, we divide the multitask features into three parts: [Z_1_], [Z_2_], and [Z_3_, Z_4_, . . ., Z_T+2_]. [Z_1_] is fed into the Category-Decoder to predict the primary category of natural images; [Z_2_] is input to the Label-Decoder to predict multiple labels of natural images; and [Z_3_, Z_4_, . . ., Z_T+2_] is input to the Text-Decoder to generate textual descriptions of natural images.

### (3) Category-Decoder

The Category-Decoder is a classifier, whose responsibility is to predict the primary category from perceptual activities. We use a common Multi-layer Perceptron (MLP) as the structure for the Category-Decoder. In Fig. 5d, the MLP employing two hidden layers for classifying the multitask feature [Z_1_]. To enhance model generalization and stability, each neuron within the hidden layers utilizes the Leaky-ReLU (0.2) activation function and Layer Normalization. The output layer of the Category-Decoder comprises 12 neurons, each representing a distinct category. To alleviate the effects of saturation, we applied the Softmax activation function to these neurons, enabling the acquisition of a probability distribution for predicting categories. Leveraging the Category-Decoder, we could compute the predicted category (C_pred_) and the category loss (L_CTG_). The specific formulas are presented below:

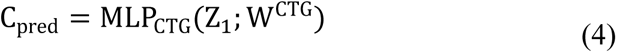

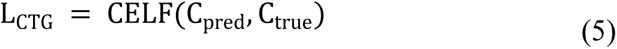

where MLP_CTG_(⋅) and W^CTG^ denote the Category-Decoder and its corresponding parameter set, respectively; [Z_1_] stands for the first part of the multitask features extracted from the Multitask-Encoder; CELF(⋅) stands for the Cross Entropy Loss Function; C_pred_ and C_true_ denote the predicted category and the actual category, respectively.

### (4) Label-Decoder

To predict multiple labels of natural images from visual activities, we employed the Label-Decoder depicted in Fig. 5e to transform the multitask feature Z_2_ into a probability distribution for 80 labels. Similar to the Category-Decoder, the Label-Decoder was also an MLP comprising two hidden layers, utilizing the Leaky-ReLU (0.2) activation function and Layer Normalization for enhanced stability. Likewise, to mitigate saturation effects, the Softmax activation function for non-linear transformation was employed in the final layer of the Label-Decoder, yielding a probability distribution for the 80 labels. Specifically, given [Z_2_] from the Multitask-Encoder, the predicted labels (S_pred_) for natural images can be obtained via the Label-Decoder. The explicit formulas for predicting labels and their associated loss are detailed below:

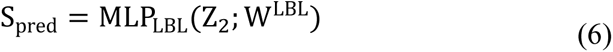

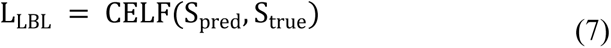

where MLP_LBL_(⋅) and W^LBL^ denote the Label-Decoder and its corresponding parameter set, respectively; Z_2_ stands for the second part of the multitask features extracted from the Multitask-Encoder; CELF(⋅) stands for the Cross Entropy Loss Function; S_pred_ and S_true_ denote the predicted labels and the actual labels, respectively.

### (5) Text-Decoder

The Text-Decoder aims to generate text describing the stimulus image from perceptual activities. In Fig. 5f, the task of the Text-Decoder was to convert multi-task features [Z_3_, Z_4_, . . ., Z_T+2_] into descriptive textual (D_pred_). This implementation involved a detailed analysis and sophisticated encoding of the features, allowing the transformation of visual information into understandable text. To capture the underlying linguistic information in brain activity, we constructed the Text-Decoder using a decoder of Transformer. Its output comprises predictions for textual descriptions (D_pred_), and the specific implementation process is outlined as follows:

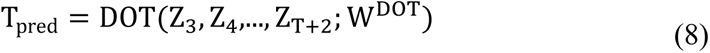

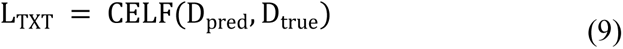

where DOT(⋅) and W^DȮT^ denote the Text-Decoder and its corresponding parameter set, respectively; [Z_3_, Z_4_, . . ., Z_T+2_] represent the third part of the multitask features obtained from Multitask-Encoder; CELF(⋅) stands for the Cross Entropy Loss Function; D_pred_ and D_true_ denote the predicted text and the actual text, respectively.

### Multi-task decoding on the VLDM

We used the NSD dataset to train and test our model. During the training phase, we employed the Adam optimizer^47^ to minimize the total loss (L_CTG_ + L_LBL_ + L_TXT_) and obtain an optimized parameter configuration. Throughout the training, we conducted 300 epochs, utilizing a learning rate of 0.001 and a batch size of 128. It’s important to note that the testing results for category decoding and label decoding were simultaneously generated during the training process. If there was no observable improvement in test performance over consecutive iterations, we would terminate the training. The entire computational process was executed using 2 NVIDIA A100 GPUs, each equipped with 40 GB of memory.

### (1) Category decoding

In the multitask decoding approach, category decoding is identified as the primary task since it provides essential information about how the brain perceives natural images. In this study, subjects viewed natural images encompassing 12 different categories (person, vehicle, outdoor, animal, accessory, sports, kitchen, food, furniture, electronic, appliance, and indoor) (depicted in Fig. 2a), and fMRI captured the response activities of these images across 10 visual cortex regions (V1, V2, V3, OFA, PPA, OPA, VWFA, FBA, FFA, and EBA). These visual activities were transformed into multitask features by the Visual-Encoder and Multitask-Encoder. Subsequently, the first segment of these multitask features was fed into the category decoder to obtain the category distribution of the natural images. We employed classification accuracy as an evaluation metric, quantitatively analyzing and measuring the accuracy of correctly categorizing viewed natural images into the 12 distinct categories. First, to compare the classification decoding performance across different cortical regions, we conducted multitask visual decoding separately on 10 distinct visual areas (V1, V2, V3, OFA, PPA, OPA, VWFA, FBA, FFA, and EBA) and 3 combined visual areas (LVC, HVC, and VC). Second, to gain insight into the intricate relationships of the 12 categories within brain representations, we performed a 2D visualization of the category distribution features output by the classification decoder with the t-distributed Stochastic Neighbor Embedding (t-SNE) Technique. This technique effectively retains spatial structural information from the original distribution features, enabling a clear visualization of the distribution patterns of different categories on a two-dimensional plane^48,49^. Initially, for the 2,770 test samples, we extracted the output of the classification decoder (category distribution features) and employed the t-SNE algorithm for processing. Throughout this process, the category distribution features of each test sample were mapped to a 2D coordinate, representing the position of that sample. Each sample was assigned to one of our designated 12 categories, distinguished using different colors for visualization. Third, to further understand the confusion among the 12 categories, we computed the confusion matrix. This confusion matrix is crucial for analyzing the model’s performance.

### (2) Label decoding

Unlike category decoding, the label decoding task not only predicts the primary labels (categories) within the images but also forecasts secondary labels, such as foreground or background labels. The process of label decoding includes: 1) acquiring multi-task features based on Visual-Encoder and Multitask-Encoder, similar to category decoding; 2) the Label-Decoder receives and processes the second part of multi-task features to obtain a probability distribution for the 80 labels; 3) quantitatively assessing the performance of label decoding using the mAP (Mean Average Precision) metric^50^. To investigate the performance differences between different types of information, we categorized 80 labels into two pairs of attributes such as “moving attributes” vs “static attributes” and “natural attributes” vs “artificial attributes”.

### (3) Text decoding

Text decoding is more complex than label decoding. This is because it not only requires the proposed model to predict labels in the stimulus image but also to string together these labels to form a sentence describing the entire content of the stimulus image. To achieve this goal, we associated brain visual activity with text describing the stimulus image and trained a text decoder based on an attention mechanism. This decoder can first extract multi-task features from the output of the multi-task encoder, and generate one word at a time, gradually composing a complete descriptive sentence. At each step of the generation process, the model determines a candidate list for the next word based on the words generated so far and the multi-task features. To quantitatively assess the accuracy of language decoding, we employed a series of evaluation metrics, including BLEU, CIDEr, ROUGE, WCS, GCS, and FTCS, to measure the similarity between the decoded sentences and the target sentences. Among these metrics, WCS, GCS, and FTCS utilize Word2vec, Glove, and FastText embeddings, respectively, to obtain sentence vectors by computing average word vectors. Then, the cosine similarity between the sentence vectors of the decoded sentence and the corresponding target sentence were calculated. The reason for selecting WCS, GCS, and FTCS as evaluation metrics is that they can accurately measure the semantic similarity between decoded sentences and target sentences. Further, to investigate preferences within the visual cortex for language decoding, we assessed ten distinct regions (including V1, V2, V3, EBA, FBA, FFA, OFA, OPA, PPA, and VWFA) along with three combined areas (LVC, HVC, and VC) for their decoding accuracy using Word2vec. We chose Word2vec due to its demonstrated effectiveness in assessing language relevance and measuring semantic similarity^20^.

## Acknowledgements

This work was supported by STI 2030-Major Projects 2022ZD0208900, the National Natural Science Foundation of China (Nos. 62333003, 62036003, 82121003, 62276051); Medical-Engineering Cooperation Funds from University of Electronic Science and Technology of China (ZYGX2021YGLH201); Natural Science Foundation of Sichuan Province (2023NSFSC0640); the Second Round Research Project of Chongqing First-class Discipline Foreign Language and Literature (Grant No. SISUWYJY202305).

## Data availability

The NSD dataset was provided by Allen and colleagues^28^. The corresponding author can supply all other datasets upon a reasonable request.

## Code availability

The full set of source code necessary for the processing of datasets, along with the training and evaluation of the models and methods discussed, can be accessed at https://github.com/hw654833612/Visual-Language-Decoding.

## Author contributions

The project was led by H.F.C. and H.M.Y. W.H., Y. H. Z., J. W., H. X. L., J. P. L., and Y. C. Z. took care of the data curation. The training/evaluation pipeline, including model training and hyperparameter search, was built and handled by W.H., F.Q., W. R., and S. Z. W. W.H., P.F.Y., Y.T. and D.W.W. provided in depth data and results analysis. The present paper was written by W.H., K.W.C., J. Y. S., Q. L., B. Z., and H.J.L.

## Supplementary Materials for “From Sight to Insight: A Multi-task Approach with the Visual Language Decoding Model”

**Supplementary Table 1.**
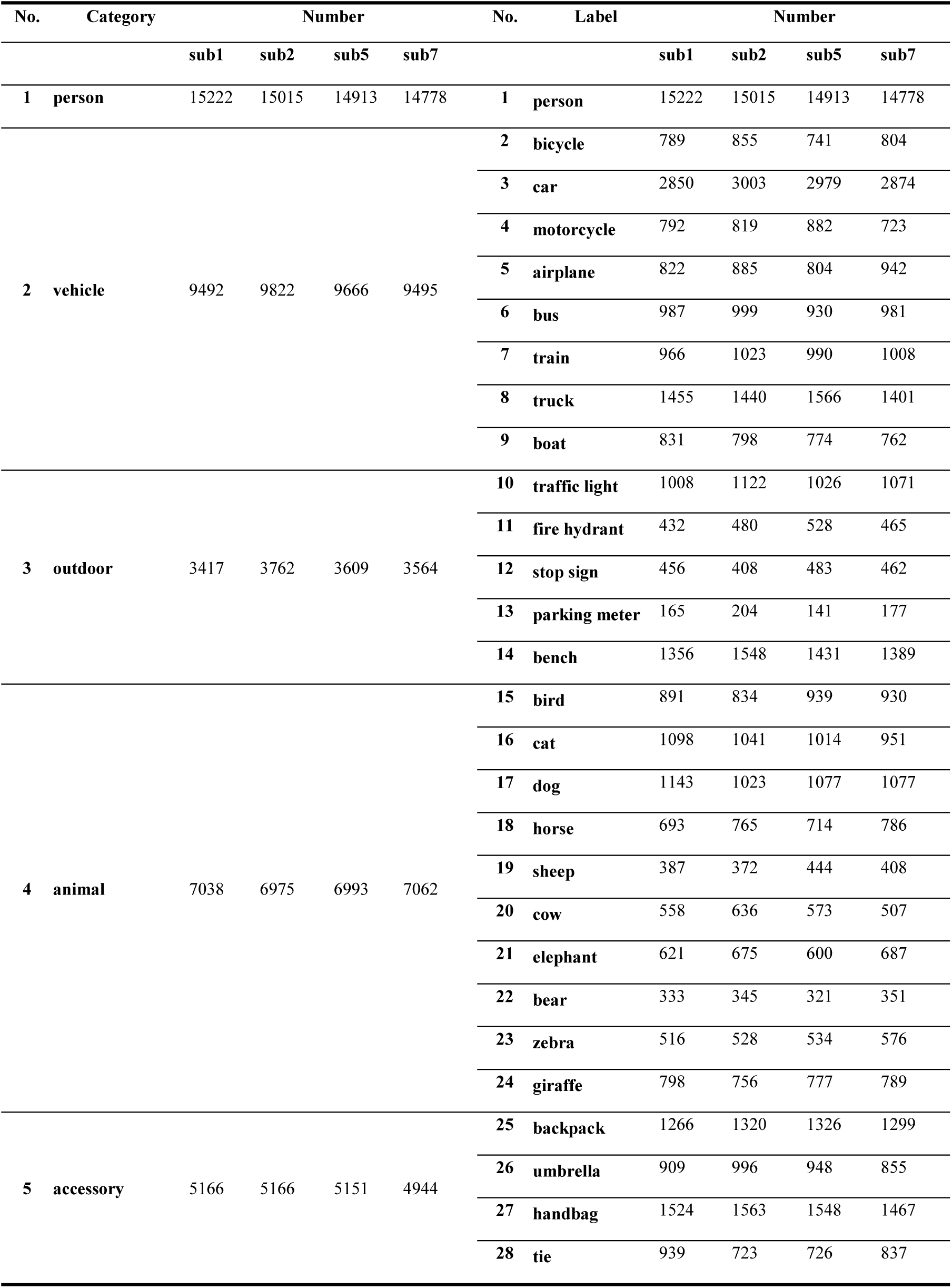
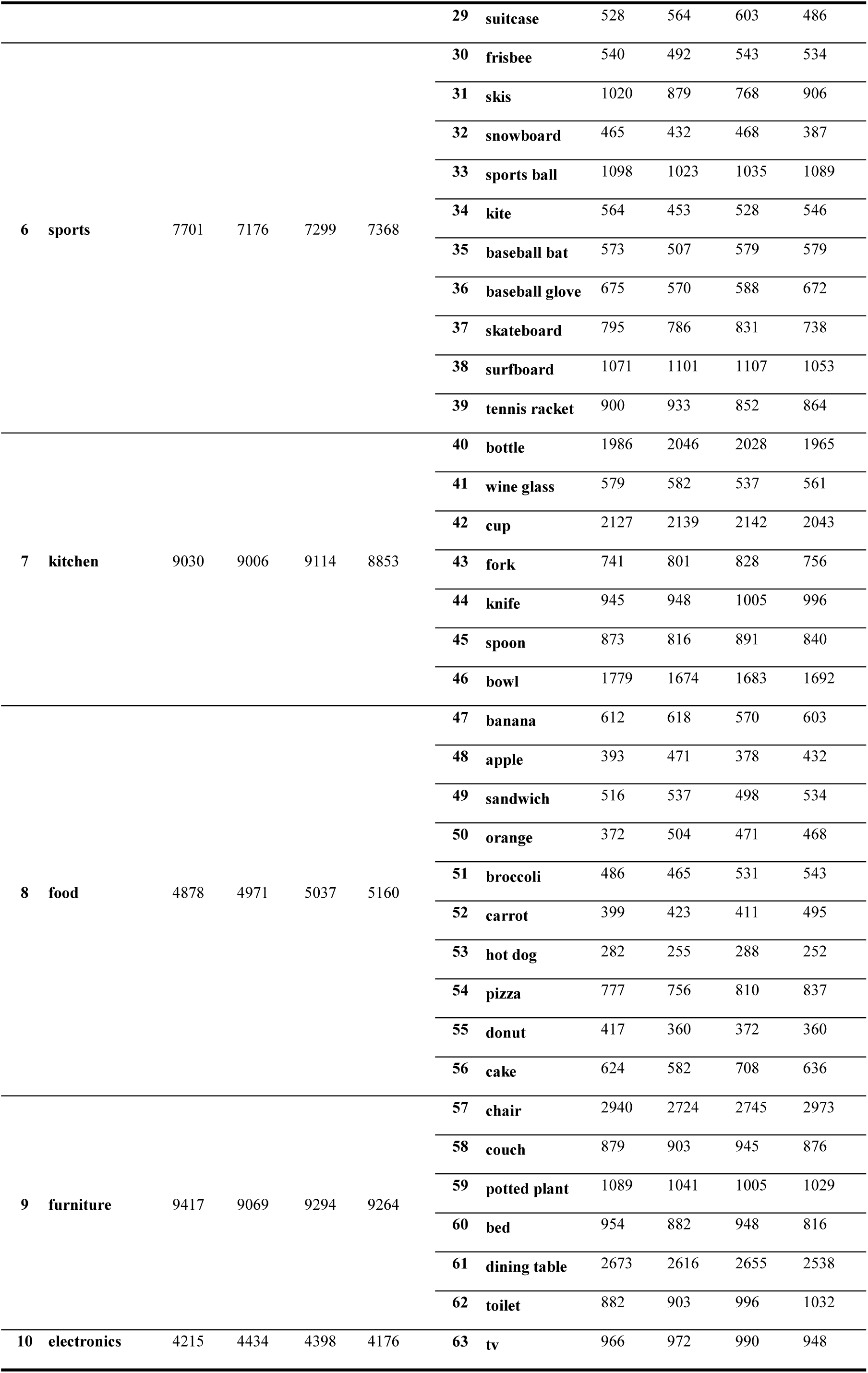
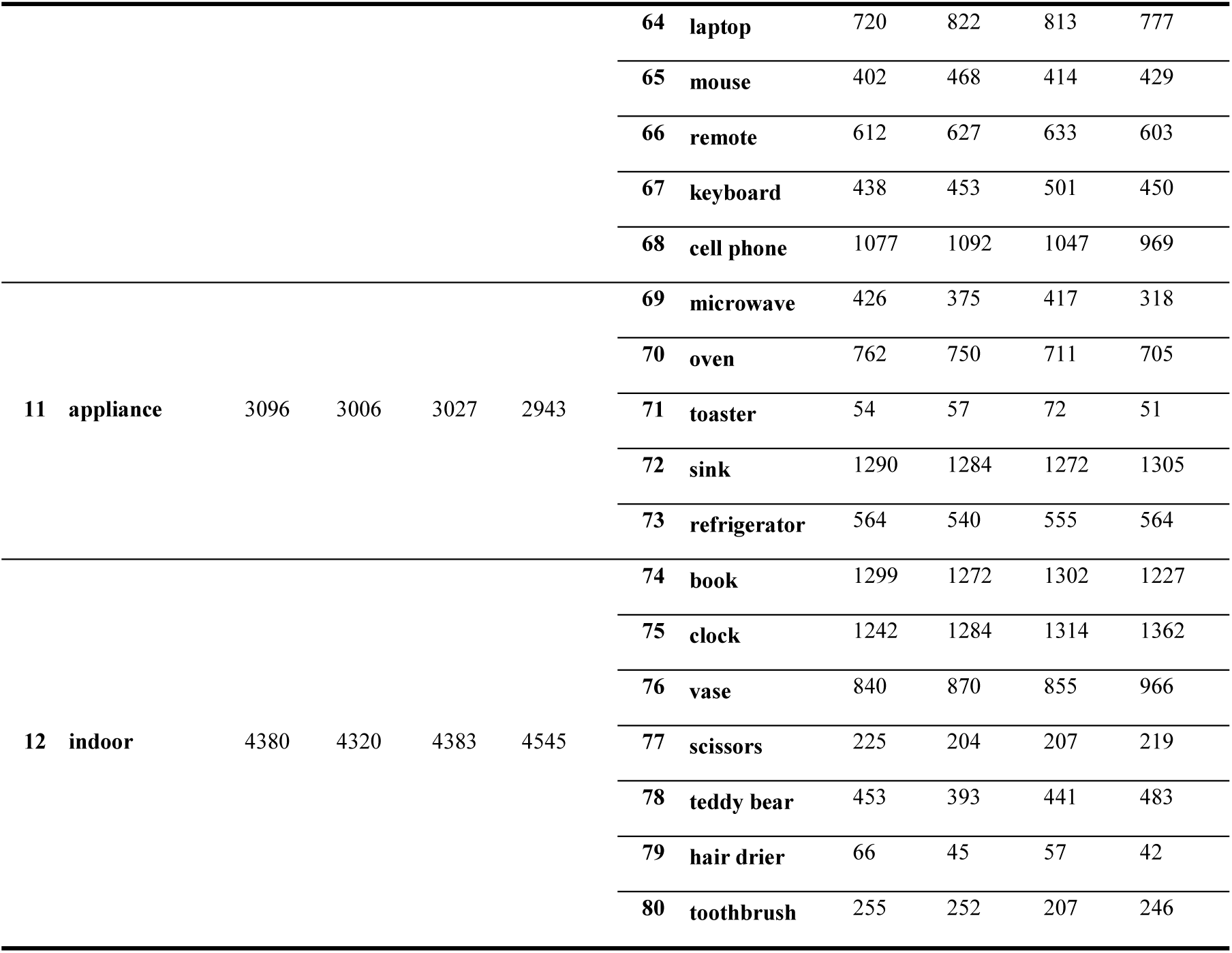
Statistical quantities of elements for ‘primary categories’ and ‘multi labels’ in NSD.

**Supplementary Fig. 1.**
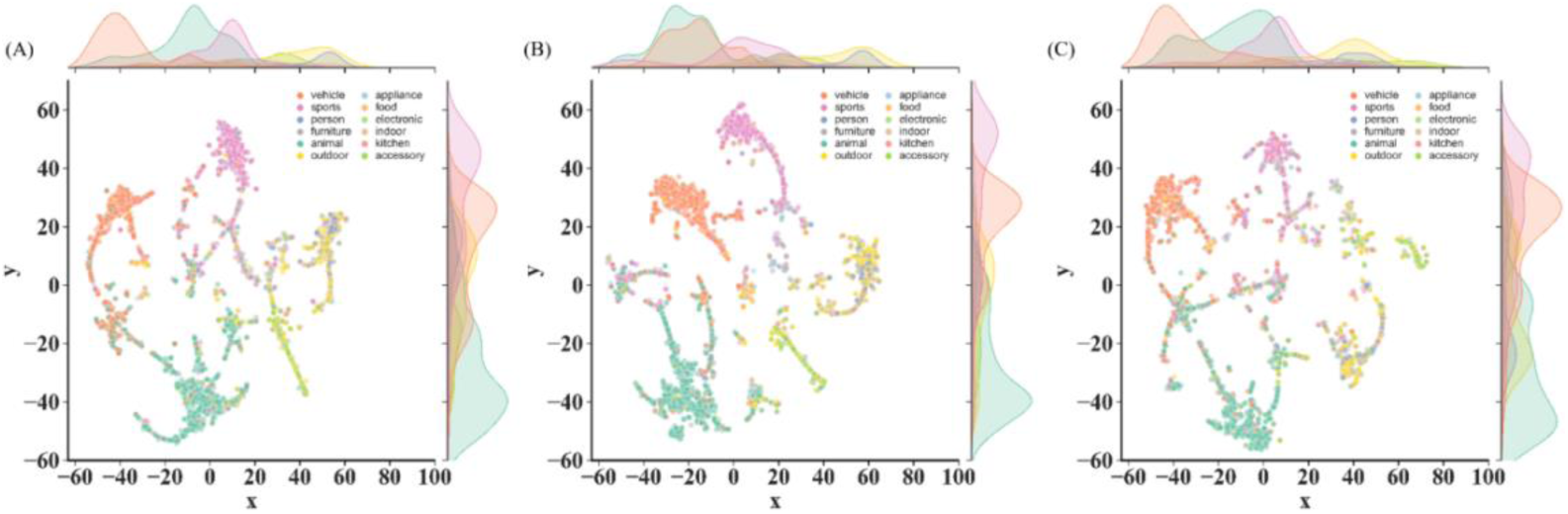
The output features of Category-Decoder for Subject 2, Subject 5 and Subject 7 are shown from left to right, which were downscaled by T-SNE to a two-dimensional space to generate a visualization map of the 12 categories.

**Supplementary Fig. 2.**
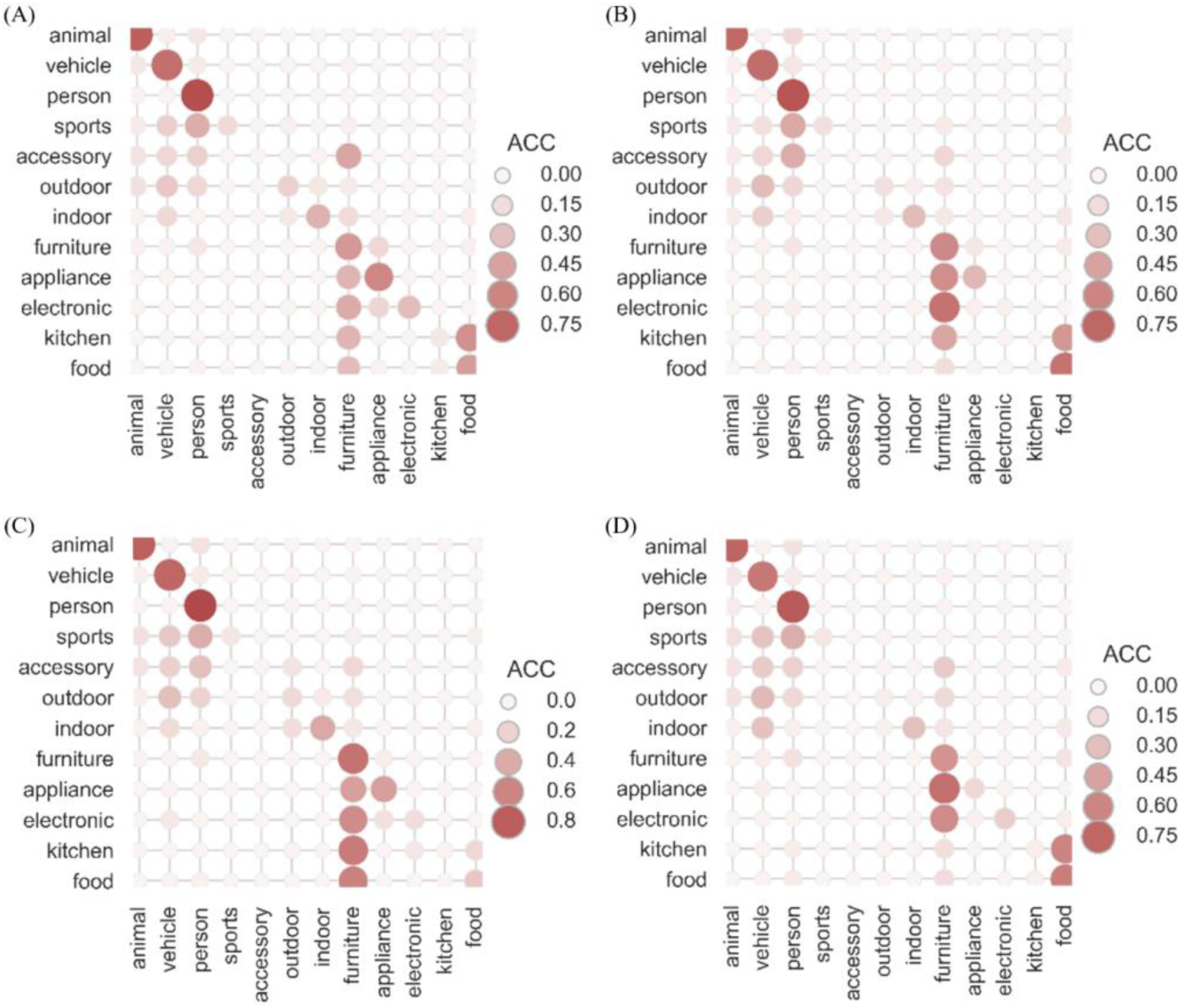
The four subfigures show the confusion matrices for the 12 categories obtained from the response in the VC area corresponding to Subject 1, Subject 2, Subject 5 and Subject 7.

**Supplementary Fig. 3.**
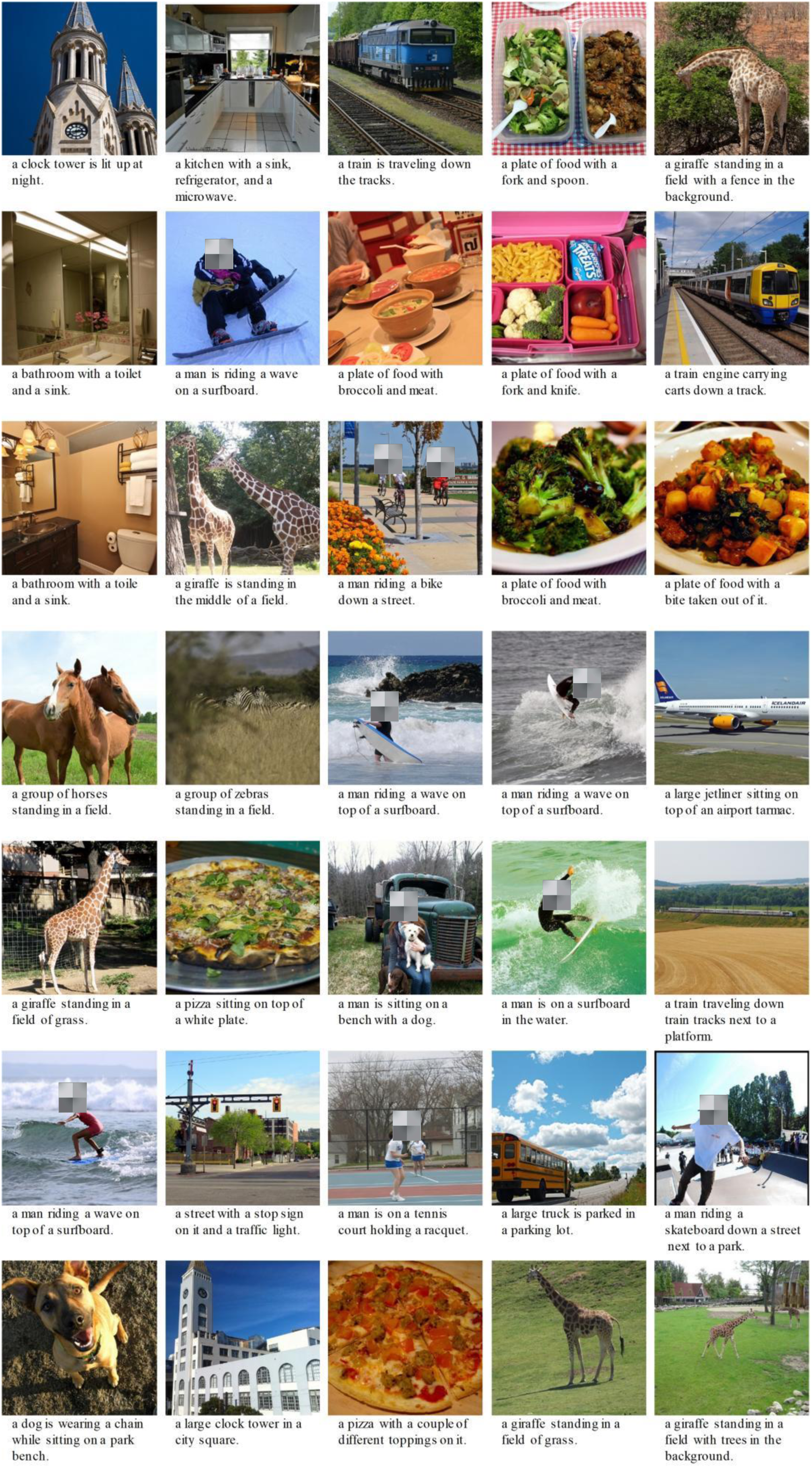
Each image is derived from the test set, accompanied by the text generated by the Text-Decoder describing the stimulus image based on brain activity.

**Supplementary Fig. 4.**
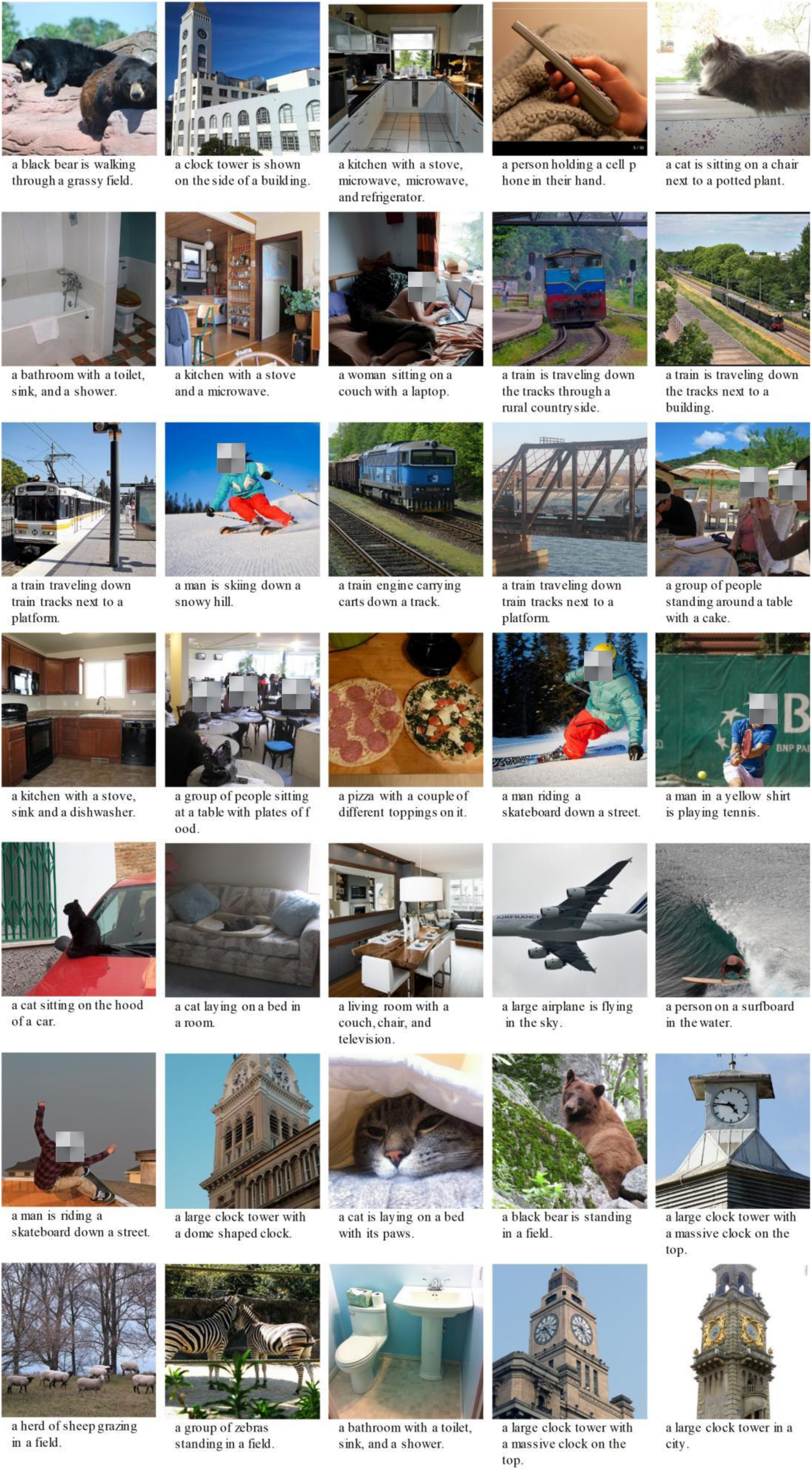
Each image is derived from the test set, accompanied by the text generated by the Text-Decoder describing the stimulus image based on brain activity.

**Supplementary Fig. 5.**
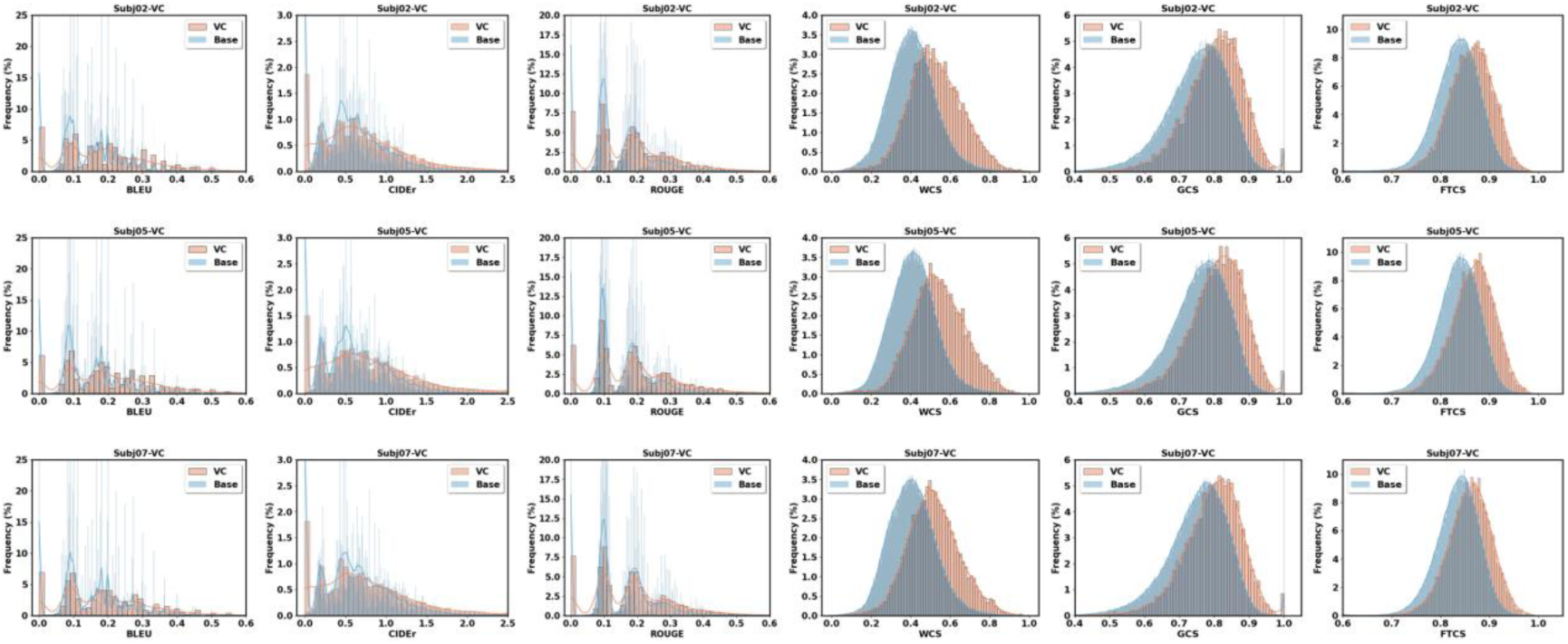
The upper, middle and lower rows show the frequency distributions of six evaluation metrics (BLEU, CIDEr, ROUGE, WCS, GCS, and FTCS) obtained by Subject 2, Subject 5, and Subject 7, respectively, in the VC response. In each subplot, the orange distribution represents the evaluation values between the decoded text and the standardized text, while the blue distribution, which serves as the baseline, indicates the distribution of evaluation values between the decoded text and all the texts in the test set.

